# The respiratory enzyme complex Rnf is vital for metabolic adaptation and virulence in *Fusobacterium nucleatum*

**DOI:** 10.1101/2023.06.13.544113

**Authors:** Timmie A. Britton, Chenggang Wu, Yi-Wei Chen, Dana Franklin, Yimin Chen, Martha I. Camacho, Truc T. Luong, Asis Das, Hung Ton-That

## Abstract

A prominent oral commensal and opportunistic pathogen, *Fusobacterium nucleatum* can traverse to extra-oral sites such as placenta and colon, promoting adverse pregnancy outcomes and colorectal cancer, respectively. How this anaerobe sustains many metabolically changing environments enabling its virulence potential remains unclear. Informed by our genome-wide transposon mutagenesis, we report here that the highly conserved Rnf complex, encoded by the *rnfCDGEAB* gene cluster, is key to fusobacterial metabolic adaptation and virulence. Genetic disruption of the Rnf complex via non-polar, in-frame deletion of *rnfC* (Δ*rnfC*) abrogates polymicrobial interaction (or coaggregation) associated with adhesin RadD and biofilm formation. The defect in coaggregation is not due to reduced cell surface of RadD, but rather an increased level of extracellular lysine, which binds RadD and inhibits coaggregation. Indeed, removal of extracellular lysine via washing Δ*rnfC* cells restores coaggregation, while addition of lysine inhibits this process. These phenotypes mirror that of a mutant (Δ*kamA*) that fails to metabolize extracellular lysine. Strikingly, the Δ*rnfC* mutant is defective in ATP production, cell growth, cell morphology, and expression of the enzyme MegL that produces hydrogen sulfide from cysteine. Targeted metabolic profiling demonstrated that catabolism of many amino acids, including histidine and lysine, is altered in Δ*rnfC* cells, thereby reducing production of ATP and metabolites including H_2_S and butyrate. Most importantly, we show that the Δ*rnfC* mutant is severely attenuated in a mouse model of preterm birth. The indispensable function of Rnf complex in fusobacterial pathogenesis via modulation of bacterial metabolism makes it an attractive target for developing therapeutic intervention.

**IMPORTANCE:** Although viewed as an oral commensal, the Gram-negative *F. nucleatum* is an opportunistic pathogen that can spread to extra-oral sites such as placenta and colon, promoting adverse pregnancy outcomes and colorectal cancer, respectively. How this anaerobe sustains various metabolically changing environments enabling its virulence potential remains unclear. We demonstrate here that the highly conserved Rnf complex is key to fusobacterial metabolic adaptation and virulence. Genetic disruption of this Rnf complex causes global defects in polymicrobial interaction, biofilm formation, cell growth and morphology, H_2_S production, and ATP synthesis. Targeted metabolomic profiling demonstrates that the loss of this respiratory enzyme significantly diminishes catabolism of numerous amino acids, which negatively impacts fusobacterial virulence as tested in a preterm birth model in mice.

## INTRODUCTION

A prominent member of the human oral microbiota that is known to harbor over 700 bacterial and fungal species (1, 2), the Gram-negative anaerobe *Fusobacterium nucleatum* plays an integral role in oral biofilm development and dental plaque formation, by virtue of its inherent capacity to adhere to diverse microbial species, notably the *Actinomyces* spp., *Streptococcus* spp., *Porphyromonas gingivalis*, and *Candida albicans* (3-12). Beyond commensalism, *F. nucleatum* is also an opportunistic pathogen that can induce preterm birth, promote tumor growth and metastatic progression of breast cancer cells, proliferation and migration of pancreatic cancer cells, and colorectal cancer (CRC), and noted for its ability to spread from the oral cavity and colonize many extra-oral sites, including placental, breast, pancreatic, and colorectal tissues (13-17). To date, a number of adhesins have been shown to mediate host-pathogen interactions. These include Fap2 and FadA, with the former aiding fusobacterial binding to tumor- and placenta-expressed Gal-GalNAc (18, 19), while the latter promoting placental colonization and CRC progression (17, 20-22). The other known adhesin RadD is a major fusobacterial virulence factor that not only mediates polymicrobial interaction (or coaggregation), a process that is inhibited by arginine and lysine (23, 24), but it is also critical for adverse pregnancy outcomes in a mouse model of preterm birth (24).

Recently, a genome-wide Tn5 transposon mutagenesis screen for identifying additional coaggregation factors revealed that the genetic disruption of a lysine metabolic pathway (LMP), e.g. deletion of *kamA* and *kamD* genes, blocks coaggregation through the excess accumulation of extracellular lysine, which binds and inhibits RadD (24). The subsequent discovery that the *kamA* deletion mutant is significantly attenuated in inducing preterm birth in mice (24) led to the realization that amino acid metabolism might play a key role in fusobacterial virulence. Consistent with this, targeted genetic analyses showed that a mutant, Δ*megL*, defective in cysteine/methionine metabolism leading to decreased hydrogen sulfide (H_2_S) production, is also attenuated in virulence in the mouse model of preterm birth (25). Yet another set of genes whose disruption by Tn5 insertion mutagenesis obliterated both coaggregation and biofilm development by fusobacteria encode a putative respiratory enzyme known as the *<u>R</u>hodobacter* <u>n</u>itrogen-fixation (Rnf) complex (26).

Originally identified in *Rhodobacter capsulatus* (27), the Rnf complex is a highly conserved and evolutionarily ancient membrane-bound ferredoxin:NAD+ oxidoreductase that is thought to couple reversible electron transfer from reduced ferredoxin (fd_red_) to NAD^+^ with the establishment of an ion-motive force (IMF), enabling substrate import and/or ATP biosynthesis (27-29). Found in over 150 bacterial genomes and 2 archaeal genomes, with high occurrence in anaerobes (28), the Rnf complex has been demonstrated to be a multifunctional respiratory enzyme that acts as a versatile metabolic exchange center for N_2_-fixation (27), carbon dioxide-fixation (30), metabolism of low-energy substrates, such as ethanol and lactate (31-33), gene regulation to some extent (34, 35), and gut colonization in mice (36). Importantly, whether the Rnf complex contributes to bacterial virulence remains to be explored. Here, we show that *F. nucleatum* encodes a functional *rnf* locus, *rnfC*-*rnfD*-*rnfG*-*rnfE*-*rnfA*-*rnfB*, and that genetic disruption of the Rnf complex, via deletion of *rnfC* or *rnfD*, causes severe defects in many virulence traits of *F. nucleatum*, including coaggregation, biofilm formation, H_2_S production, and ATP production, in addition to altering cell morphology and growth. Intriguingly, while the defect in coaggregation is mechanistically linked to a failure of lysine catabolism, leading to an increased level of extracellular lysine that inhibits RadD-mediated coaggregation, the defects in other traits are attributed to global reduction of amino acid metabolism as determined by targeted metabolomic analysis. Most significantly, the *rnfC* mutant is severely impaired in inducing preterm birth in mice. This study establishes that the Rnf complex is a central component in *F. nucleatum* metabolism that broadly impacts bacterial virulence.

## RESULTS

### Transposon mutagenesis reveals the involvement of the *F. nucleatum* Rnf complex in biofilm formation, coaggregation, and H_2_S-production.

In our previous genome-wide screens that identified *F. nucleatum* mutants defective in biofilm formation and coaggregation, multiple subunit-encoding genes for the predicted Rnf complex were frequently targeted by Tn5 transposon insertion (24, 26) (Fig. 1A). To establish the potential multifunctional role of the Rnf complex, we further characterized these mutants in standard biofilm and coaggregation assays (24, 26). To cultivate monospecies biofilms, normalized fusobacterial cultures of different strains were seeded into sterile multi-well plates, anaerobically cultured at 37°C for 48 h, and the resultant biofilms were then stained with crystal violet. Compared to the wild-type strain (ATCC 23726), each of the four tested mutant strains were defective in biofilm formation (Fig. 1B). To examine whether these mutants are also deficient in coaggregation, normalized, unwashed fusobacterial cells were mixed and incubated with *S. oralis* or *A. oris* in a standard assay, followed by imaging (see Methods). The tested Tn5 mutants also failed to co-aggregate with these two oral co-colonizers at a level similar to that of a *radD* mutant used as a negative control (Fig. 1C).

**Figure 1:**
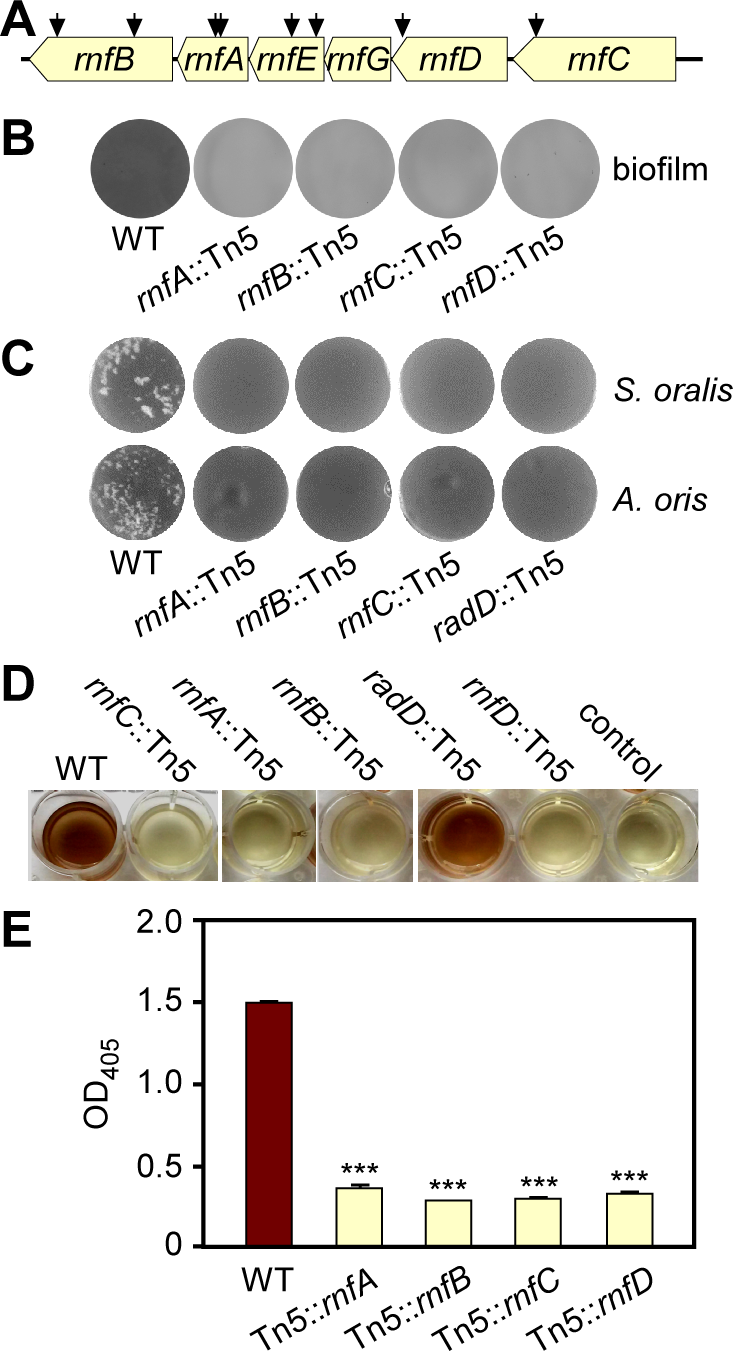
Transposon mutagenesis reveals the involvement of an Rnf complex in biofilm formation, coaggregation, and H_2_S production. **(A)** Shown is the *rnf* gene locus in *F. nucleatum*, with arrows indicating the Tn5 insertion sites of previously identified Tn5 mutants (24, 26). **(B)** Mono-species biofilms of the parent (WT) and Tn5 *rnf* mutant strains were cultivated in multi-well plates, stained with 1% crystal violet, and imaged. **(C)** Indicated fusobacterial strains were examined for their adherence to *S. oralis* and *A. oris* by a coaggregation assay. A *radD*::Tn5 mutant was used as a negative control. **(D-E)** Production of H_2_S from indicated strains was determined by the bismuth sulfide method (D) and quantified by absorbance measurement at 405 nm (E). Cell-free samples were used as a negative control. All results were obtained from three independent experiments performed in triplicate.

Unexpectedly, we noticed that all of the *rnf* mutants lacked the rotten-egg odor characteristically produced by wild-type fusobacteria, suggestive of severe deficiency in hydrogen sulfide production by these mutants. Indeed, in a bismuth sulfide assay, in which bismuth trichloride reacts with H_2_S to yield precipitation of brown bismuth sulfide, all *rnf*::Tn5 mutants showed a significantly reduced level of hydrogen sulfide relative to the wild-type strain (Fig. 1D-1E). Overall, these results point to a multifaceted role of the Rnf complex in many cellular processes in *F. nucleatum*.

### Genetic deletion of *rnfC* disrupts RadD-mediated coaggregation via blockage of lysine catabolism

A Tn5 insertion can potentially cause polar effects on transcription of downstream genes. Although this is unlikely the case here (Fig. S1A), we decided to generate non-polar, in-frame deletions of *rnf* genes utilizing our published protocols (24, 26, 37). Since all Tn5 mutants displayed similar phenotypes, we chose *rnfC* for further characterizations, as RnfC is predicted to be a peripheral membrane protein, enabling generation of polyclonal antibodies against a recombinant RnfC protein (Fig. S1B). To examine how the Rnf complex mediates polymicrobial interaction, this *rnfC* mutant (Δ*rnfC*) and other strains were subjected to the coaggregation assay, in which unwashed fusobacterial cells were mixed with *Streptococcus gordonii*, like the coaggregation assays with *S. oralis* and *A. oris* as mentioned above (see Fig. 1C). As expected, deletion of *rnfC* abrogated fusobacterial coaggregation with *S. gordonii*, as compared to the parent strain, and this defect was rescued by ectopic expression of RnfC from a plasmid (Fig. 2A; the unwashed panel).

**Figure 2:**
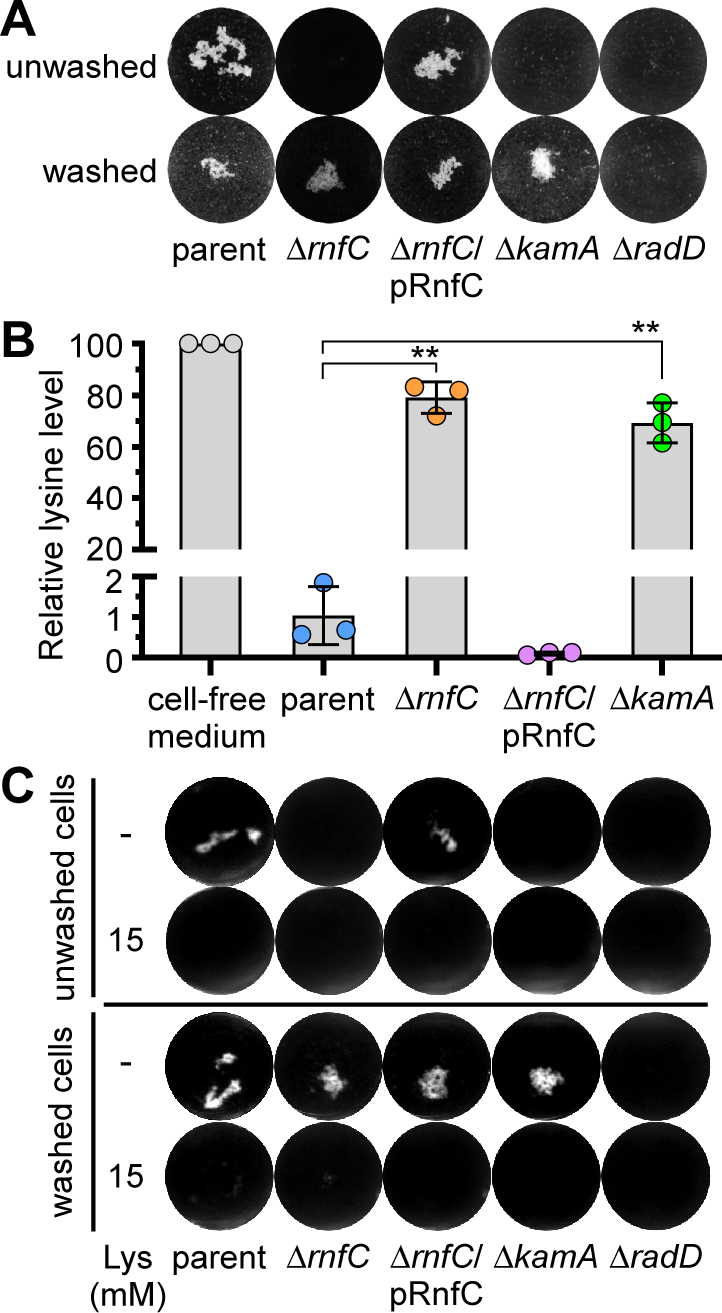
A non-polar, in-frame *rnfC* deletion mutant is defective in RadD-mediated coaggregation and lysine uptake. **(A)** The parent, *rnfC* deletion mutant, and *rnfC*-complemented strains were tested for their ability to co-aggregate with *S. gordonii.* Fusobacterial cells were washed or unwashed before being mixed in equal volumes with washed *S. gordonii* and imaged. Mutants devoid of *radD* or *kamA*, a lysine metabolic pathway gene (24), were used as references. **(B)** The relative level of lysine in the cell-free culture medium of indicated strains grown to stationary phase was determined by liquid chromatography-mass spectrometry, with the level of media without bacteria set to 100. **(C)** The same coaggregation experiment as (A) was performed with or without addition of 15 mM lysine. All results were obtained from three independent experiments performed in triplicate. Significance was calculated by a student’s t-test; *P < 0.05; **P < 0.01.

Because RadD is essential as the adhesin for fusobacterial coaggregation (23, 24), we examined if deletion of *rnfC* causes a defect in surface expression of RadD by immuno-fluorescence microscopy, wherein normalized fusobacterial cells of different strains were stained with antibodies against RadD (α-RadD), followed by counter-staining with Alexa 488-conjugated IgG. Microscopic examination presented in Fig. S2A revealed no significant defects of RadD-surface expression in the Δ*rnfC* as compared to the parent strain. Consistent with this observation, western blotting of whole cell lysates showed similar levels of RadD antigen production in all strains – parent, Δ*rnfC*, and its complement (Fig. S2B). Thus, the coaggregation defect of Δ*rnfC* is not due to reduced RadD expression.

Unexpectedly, when we utilized Δ*rnfC* cells free of the culture medium before mixing with *S. gordonii*, the mutant fusobacteria were able to adhere to oral streptococci at a level comparable to the parent strain (Fig. 2A; compare panels washed vs unwashed). This behavior of sensitivity to the culture medium mirrored that of a deletion mutant of *kamA*, which is defective in lysine catabolism and hence accumulates excess lysine in the culture medium (24). Given that lysine inhibits RadD-dependent coaggregation (23, 24), the results prompted us to examine the relative extracellular levels of lysine in these strains by using liquid chromatography-mass spectrometry as previously reported (24). Strikingly, compared to the parent strain, the extracellular level of lysine for the Δ*rnfC* mutant remained high and was quite comparable to that of the Δ*kamA* mutant (Fig. 2B). Thus, this accumulation of extracellular lysine is likely the inhibitor that blocks coaggregation by Δ*rnfC* mutant bacteria. To obtain support for this hypothesis, we re-performed the coaggregation assay with the same set of strains in the absence or presence of added 15 mM of lysine in the coaggregation mix. Regardless of whether fusobacterial cells were washed or not, the addition of lysine blocked fusobacterial coaggregation with *S. gordonii* as expected (Fig. 2C).

Since deletion of *kamA* leads to accumulation of extracellular lysine (Fig. 2B) (24), we examined the transcript level of *kamA* in the Δ*rnfC* mutant by qRT-PCR. Remarkably, expression of *kamA* was significantly reduced in the absence of *rnfC*, as compared to the parent strain, and this defect was rescued by ectopic expression of *rnfC* from a plasmid (Fig. S3). In addition, expression of *kamD*, another lysine metabolic gene in the *kamA* locus (24), mirrored that of *kamA* in these strains (Fig. S3). Altogether, the results indicate that the defect of RadD-mediated coaggregation by genetic disruption of *rnfC* is due to increased extracellular levels of lysine, likely due to the reduced expression of lysine metabolic genes by an as yet unexplored mechanism.

### Genetic deletion of *rnf*C disrupts biofilm formation, growth, cell morphology, and hydrogen sulfide production

Next, to determine how biofilm formation is dependent on the Rnf complex, we first analyzed the non-polar, in-frame *rnfC* deletion mutant in the aforementioned biofilm assay using crystal violet staining (as described in Fig. 1B above). As expected, this mutant was also unable to form monospecies biofilms compared to the parent strain, and ectopic expression of RnfC on a plasmid was sufficient to restore biofilm formation (Fig. 3A). Since the Rnf complex plays a key role in energy conservation through several bacterial metabolic pathways (27, 38, 39), we reasoned that the defects in biofilm formation by *rnf* mutants could be due to alterations with aspect to fusobacterial physiology. To investigate this further, we tested whether the Rnf complex is important in ATP biosynthesis by subjecting normalized fusobacterial cells to a microbial cell viability assay (BacTiter-Glo^TM^), which relies on the mono-oxygenation of luciferin catalyzed by luciferase in the presence of ATP. Indeed, compared to the parent strain, the Δ*rnfC* mutant drastically reduced ATP production, and complementation with an RnfC-expressing plasmid rescued the defect (Fig. 3B). Consistent with this observation, the Δ*rnfC* mutant was also severely defective in growth displaying premature growth cessation after the culture reached a suboptimal density (Fig. 3C); the mutant also displayed a morphological abnormality with short/stubby cells as examined by electron microscopy (EM) (Fig. 3D-3E).

**Figure 3:**
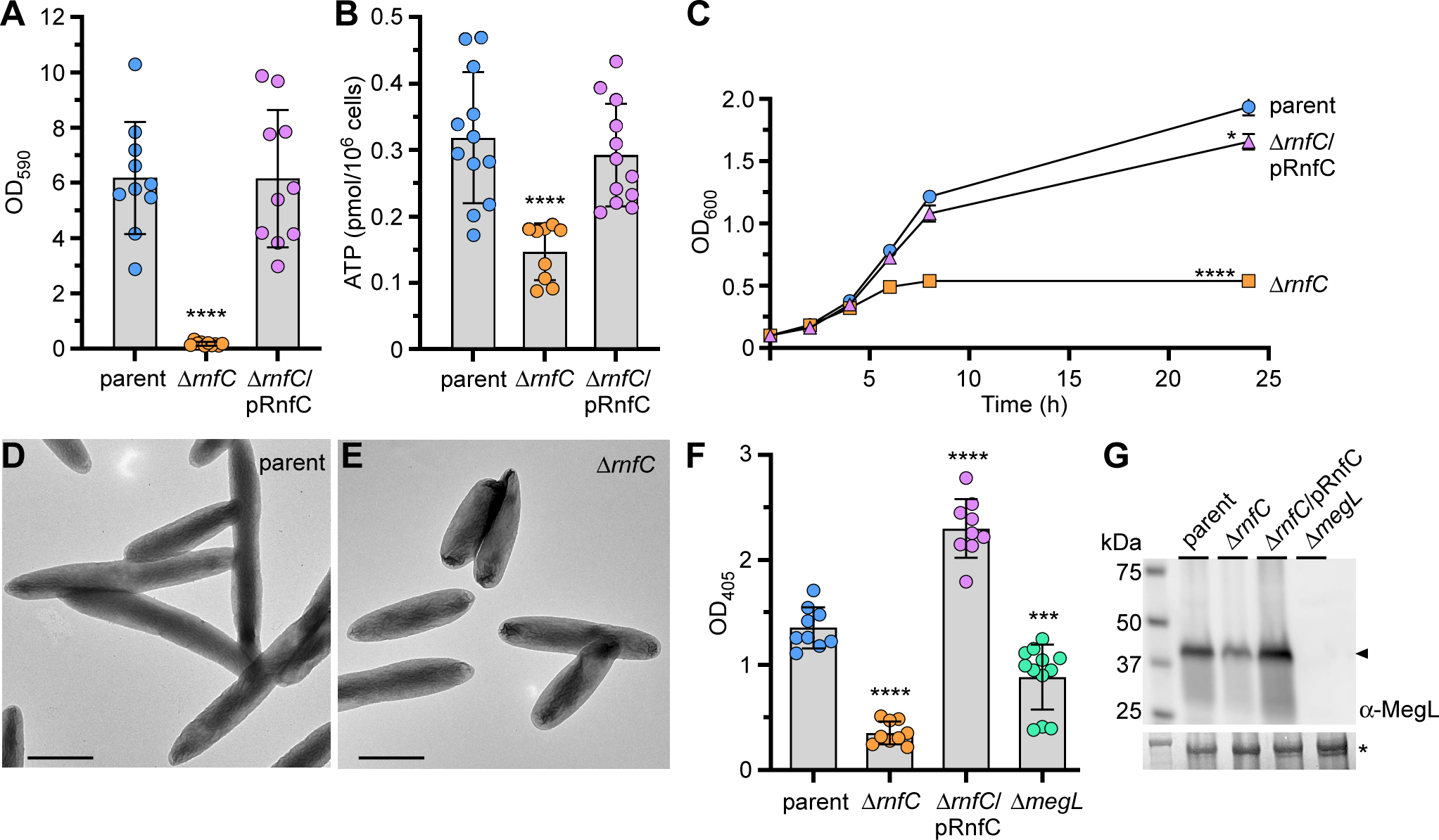
Deletion of *rnfC* causes pleiotropic defects. **(A)** 48-h grown biofilms of indicated strains were stained with 1% crystal violet and quantified by optical density measurement at 590 nm (OD_590_). **(B)** ATP production of indicated strains was assessed by a luciferase assay using a BacTiter-Glo™ kit (Promega). Mid-log phase cells were mixed with equal volumes of the BacTiter-Glo™ reagent and incubated at room temperature for 2 min before measuring luminescence on a plate reader. Normalized cell suspensions were serially diluted and plated for bacterial enumeration and subsequent normalization of relative luminescent signal. **(C)** Bacterial growth of indicated strains was determined by optical density at 600 nm at timed intervals. **(D-E)** Mid-log phase cells of parent and Δ*rnfC* mutant strains were immobilized on carbon-coated nickel grids and stained with 1% uranyl acetate prior to imaging by electron microscopy; scale bars: 1 µm. **(F)** H_2_S-production of indicated strains was determined by the bismuth sulfide assay. **(G)** Protein samples obtained from the whole-cell lysates of normalized cultures of indicated strains were subjected to immunoblotting with antibodies against MegL (α-MegL). A Coomassie Blue stained band (*) was used as a loading control. All results were obtained from three independent experiments performed in triplicate. Significance was calculated by a student’s t-test; *P < 0.05; ***P < 0.001; ****P < 0.0001.

Potentially, the physiological deficits reported above might be due to reduced metabolism of amino acids, esp. cysteine as it is one of eight important amino acids for fusobacteria (40). To test this hypothesis, we employed the aforementioned bismuth sulfide assay to measure hydrogen sulfide since it is a product of cysteine metabolism in *F. nucleatum* (25, 41-44). Consistent with the results of our Tn5 mutants, the Δ*rnfC* mutant showed a significant defect in H_2_S production, and overexpression of RnfC from a plasmid substantially enhanced this process (Fig. 3F). Because methionine-gamma lyase MegL was previously shown to be responsible for the bulk of H_2_S production from cysteine metabolism in *F. nucleatum* (25), we went on to determine the overall expression level of MegL in cells by immunoblotting of whole cell lysates with antibodies against MegL (α-MegL). Remarkably, compared with the parent and rescued strains, the Δ*rnfC* mutant expressed a significantly reduced level of MegL (Fig. 3G). This reduction of MegL enzyme corresponded to a reduced level of *megL* mRNA as determined by qRT-PCR (Fig. S3). Notably, the expression of *cysK1* and *cysK2*, which encode two additional H_2_S-producing enzymes (24), was also reduced in the absence of *rnfC* (Fig. S3).

To ascertain that the physiological defects reported above are a reflection of the loss of Rnf function, we repeated these experiments with a non-polar, in-frame *rnfD* deletion mutant (Δ*rnfD*). Like the Δ*rnfC* mutant, the Δ*rnfD* mutant showed significant defects in coaggregation, biofilm formation, cell growth, and H_2_S production, and these defects were rescued by ectopic expression of *rnfD* (Fig. S4). Altogether, these findings support our hypothesis that the biofilm formation defect caused by genetic disruption of the Rnf complex is due to gross deficiencies in ATP biosynthesis and amino acid metabolism.

### Loss of RnfC complex disrupt global *F. nucleatum* metabolism

To gain further insight into the function of the *F. nucleatum* Rnf complex in relation to the aforementioned phenotypes associated with genetic disruption of the Rnf complex, we performed metabolomic analysis of parent and Δ*rnfC* cells using ultra-performance liquid chromatography coupled to mass spectrometry (UPLC/MS). Among over 80 metabolites detected in these samples, 10 metabolites (Gln, Tyr, Ile, etc) were elevated, whereas 7 metabolites, including purine ribonucleoside precursors, were significantly depleted in the Δ*rnfC* mutant, as compared to the parent strain (Fig. 4A).

**Figure 4:**
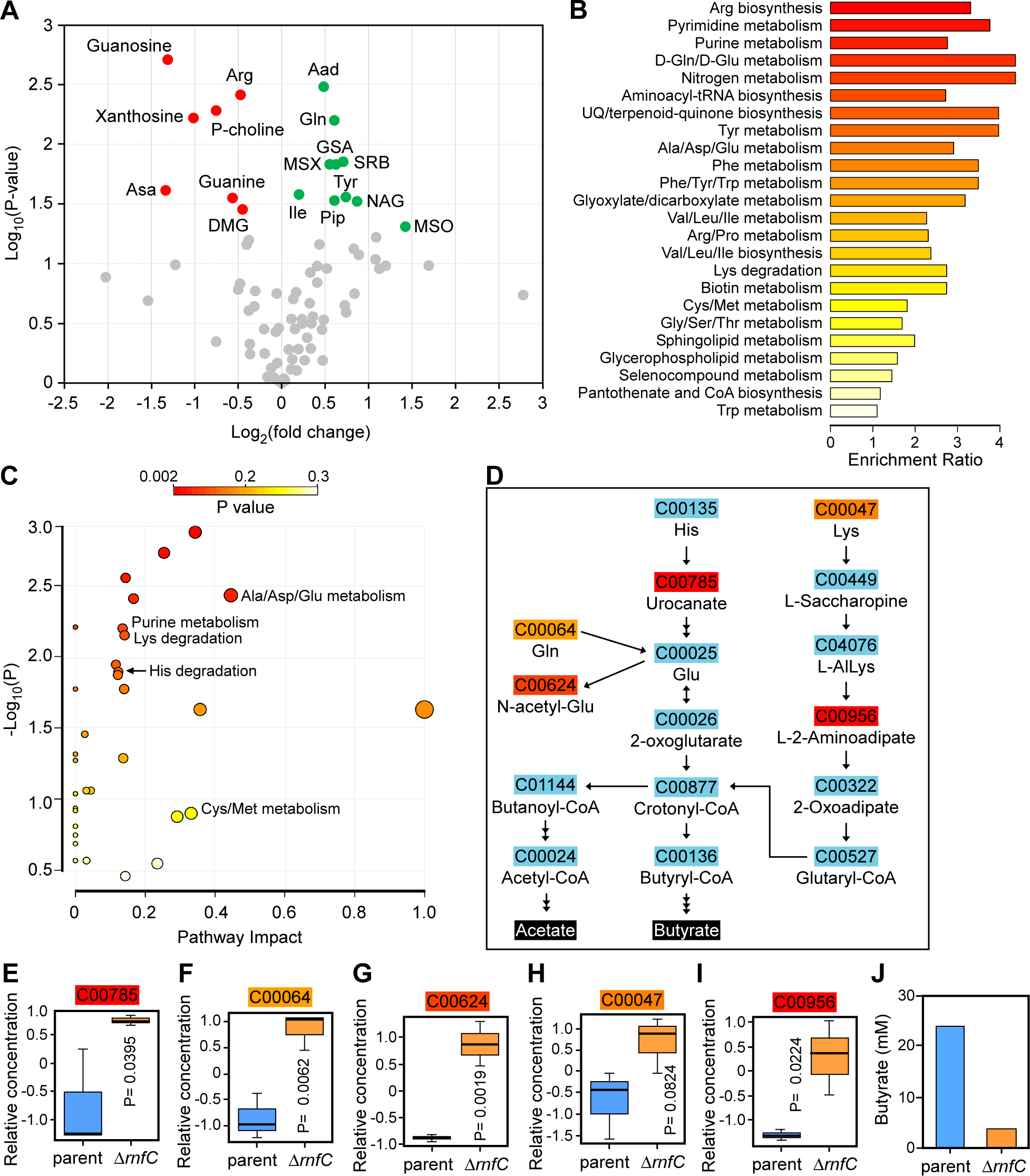
Deletion of *rnfC* disrupts amino acid metabolism. **(A)** The parent and Δ*rnfC* cells grown to mid-log phase were subjected to metabolomics analysis using liquid chromatography-mass spectrometry (LC-MS). Shown is a volcano plot of 81 differentially expressed metabolites (DEMs) between the parent and Δ*rnfC* mutant strains. Metabolites significantly depleted or enriched in the Δ*rnfC* strain, relative to the parent, are marked in red or green, respectively. **(B)** Shown is a graphical overview of quantitative enrichment analysis of DEMs between the parent and Δ*rnfC* strains generated using MetaboAnalyst 5.0. The top 24 pathway-associated metabolic sets in Δ*rnfC* compared to the parent strain were sorted based on their fold enrichment and P-values. **(C)** Pathway analysis was performed using MetaboAnalst 5.0, which combines pathway enrichment and topology analysis. A range of P-values is shown from red to yellow. **(D)** Shown are amino acid metabolic nodes in *F. nucleatum*, based on the significance level in (C), generating acetate and butyrate. **(E-I)** The relative concentrations of DEMs in the amino acid nodes shown in D between the parent and Δ*rnfC* mutant strains are presented. P-values were calculated using the Global test. **(J)** The relative level of butyrate (mM) in the overnight cultures of the parent and Δ*rnfC* strains was determined by liquid chromatography-mass spectrometry (LC-MS).

To obtain a better view of the specific metabolic pathways altered upon loss of *rnfC*, we performed a quantitative pathway enrichment and pathway topology analysis of the parent and Δ*rnfC* cell using the free, web-based software, MetaboAnalyst 5.0. In keeping with our initial findings (Fig. 4A), several pathways involved in amino acid and purine metabolism were significantly enriched in the Δ*rnfC* mutant compared to the parent strain (Fig. 4B-4C). Pathway analysis of the “Lys degradation”, “His degradation”, and “Ala/Asp/Gln metabolism” nodes in the topology analysis revealed that many amino acids are directly degraded into glutamate, which is subsequently fermented into the short-chain fatty acids, butyrate, and acetate (Fig. 4D). Significantly, the Δ*rnfC* mutant had elevated levels of urocanate, glutamine, N-acetyl-L-glutamate, lysine, and the lysine-degradation pathway intermediate, L-2-aminoadipate, relative to the parent strain (Fig. 4E-I), indicating possible disruptions in glutamate fermentation. Consistent with this result, we observed a reduced level of butyrate in this mutant as determined by high-performance liquid chromatography coupled to mass-spectrometry (HPLC/MS) (Fig. 4J). Additionally, as demonstrated by pathway analysis of the “Cys/Met metabolism” node from the topology analysis, hydrogen sulfide production is derived primarily from methionine degradation into L-homocysteine and subsequent L-cysteine degradation into pyruvate (Fig. S5A), both catalyzed by MegL (25). Correspondingly, elevated levels of methionine and slightly lower levels of the pathway intermediate, S-adenosyl-L-homocysteine, were observed in the Δ*rnfC* mutant, relative to the parent strain (Fig. S5B), suggesting the Rnf complex may also be important for methionine/cysteine metabolism. In sum, the results support a broad role of the Rnf complex in *F. nucleatum* metabolism.

### The Rnf complex is required for *Fusobacterium nucleatum* virulence

Because genetic disruption of the Rnf complex causes significant defects in many virulence traits of *F. nucleatum* (Fig. 1-3), we sought to determine whether this complex contributes to *F. nucleatum* pathogenesis. Utilizing a mouse model of pre-term birth as previously reported (24, 37), we infected via tail vein injection groups of five CF-1 pregnant mice at day 16/17 of gestation with roughly 5.0 x 10^7^ CFU of either the parent or Δ*rnfC* strain, and pup delivery was monitored (Fig. 5A). Strikingly, the Δ*rnfC* mutant was severely attenuated in virulence, with nearly 50% of pups born alive by the end-point, while the parent strain with almost no pub survival observed within the same timeframe (Fig. 5B). Clearly, the Rnf complex plays a significant role in metabolism that is central to *F. nucleatum* pathogenesis.

**Figure 5:**
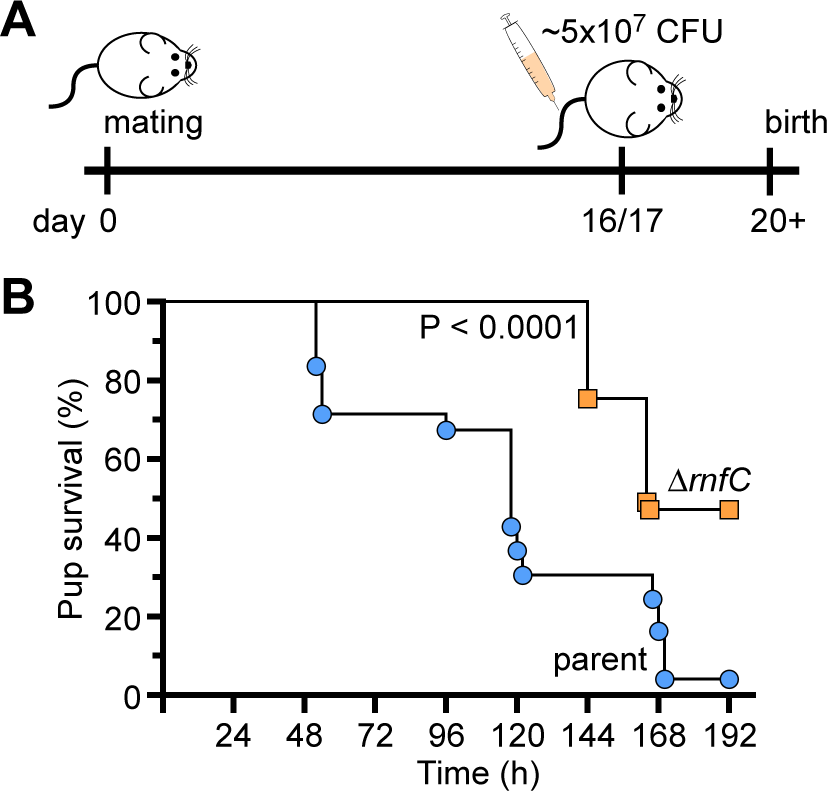
The *rnfC* mutant is attenuated in virulence. **(A-B)** Groups of pregnant CF-1 mice were infected with ∼5.0 x 10^7^ CFU/mL of the parent or Δ*rnfC* strain via tail-vein injection at day 16 or 17 of gestation. Pup survival was recorded over time. Statistical analysis was determined by Mantel-Cox testing.

## DISCUSSION

The dual life of *F. nucleatum* as a commensal in the human oral cavity and an opportunistic pathogen in extra-oral sites raises an intriguing question as to how *F. nucleatum* is able to maintain its plasticity in metabolically changing environments. Here we have demonstrated that the highly conserved multicomponent Rnf complex plays a central role in mediating metabolism of various amino acids in *F. nucleatum* (Fig. 6), which impacts many physiological traits critical for fusobacterial virulence. Specifically, various *rnf* mutants emerged in unbiased forward genetic screens in which we sought to uncover genetic factors that govern two different aspects of *F. nucleatum* pathobiology – its ability to form biofilms and to co-aggregate with certain partner co-colonizers of the oral cavity. We then showed that that targeted disruption of the Rnf complex, via deletion of *rnfC* or *rnfD*, not only causes severe defects in coaggregation, and biofilm formation but also hydrogen sulfide production, cell morphology and growth, ATP production, and induction of preterm birth (Fig. 1-5 and Fig. S4).

**Figure 6:**
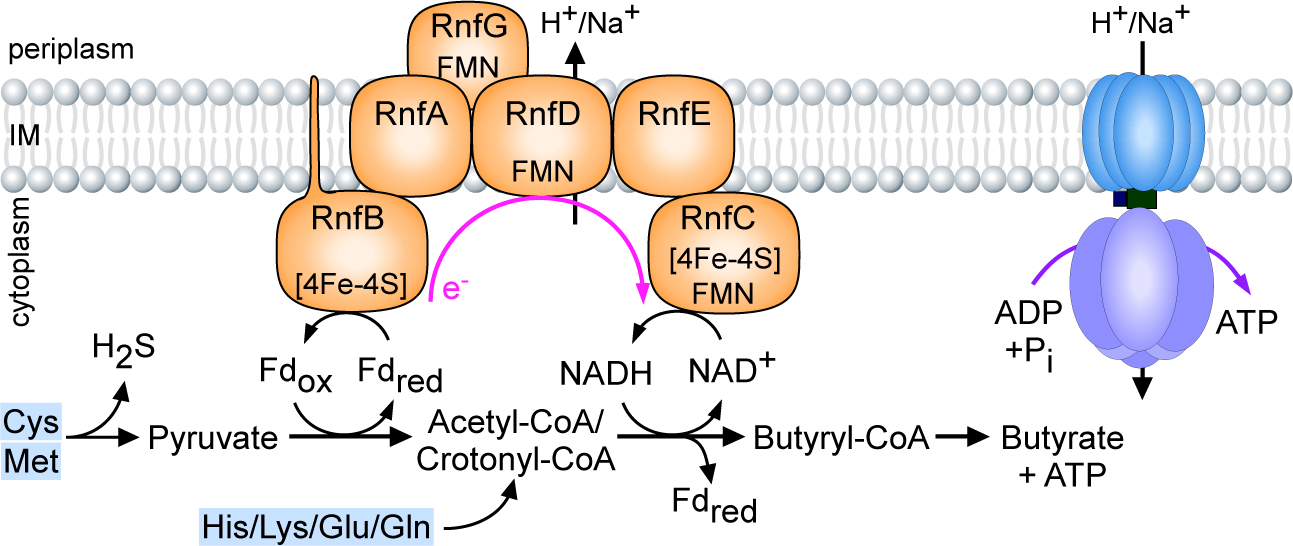
A working model of Rnf-mediated energy conservation through amino acid metabolism in *F. nucleatum*. See text for details.

The ability of fusobacteria to co-aggregate with streptococci and other partners such as *A. oris* depends on a well characterized adhesin RadD present on the fusobacterial cell surface. We have shown that the loss of Rnf complex function does not alter RadD expression or its display at the bacterial surface (Fig. S2). Rather, the defect in RadD-mediated interbacterial coaggregation appears to be a consequence of a defect in lysine metabolism mediated by the LMP, which in turn raises the level of extracellular lysine that can bind and inhibit RadD to block coaggregation (Fig. 2). Intriguingly, our results revealed that this reduced lysine metabolism parallels a decreased expression of the LMP genes, i.e. *kamA* and *kamD*, (Fig. S3). Our metabolomic profiling further revealed that the defect in amino acid metabolism is not limited to lysine alone but rather the metabolism of many additional amino acids (viz. cysteine, histidine, and glutamate) is also affected (Fig. 4 and Fig. S5). Critically, cysteine metabolism that generates hydrogen sulfide in *F. nucleatum* is mainly catalyzed by MegL, a conserved L-methionine γ-lyase (25). Logically, we expected that the defective cysteine metabolism in the Δ*rnfC* mutant would lead to reduced hydrogen sulfide production by the mutant. What we did not anticipate, however, is that the level of hydrogen sulfide production in the Δ*rnfC* mutant is significantly lower than that of the *megL* mutant (Fig. 3F), indicating the Rnf complex plays a broader role in amino acid metabolism.

Indeed, pathway and gene expression analyses demonstrated that multiple pathways leading to hydrogen sulfide production are significantly affected in the absence of *rnfC* (Fig. S5) and that expression of other cysteine metabolic genes, e.g. *cysK1* and *cysK2*, is markedly reduced (Fig. S3). Intriguingly, given CysK1 catalyzes the conversion of cysteine to hydrogen sulfide and L-lanthionine, an essential amino acid required for formation of fusobacterial peptidoglycan (45, 46), the reduced expression of *cysK1* by deletion of *rnfC* can be attributed to aberrant cell growth, short and stumpy cell size, coupled with reduced ATP production (Fig. 3). How the absence of *rnfC* reduces expression of LMP genes and *cysK1*/*cysK2* genes remains a significant puzzle. It is noteworthy that the expression of LMP genes and *megL* is controlled by the two-component transduction systems (TCSs), CarRS and ModRS, respectively (24, 25); one intriguing possibility, which remains to be tested in future studies, is that metabolic blockage by genetic disruption of the Rnf complex causes accumulation of certain metabolites and intermediates that may trigger gene expression responses from these TCSs and perhaps other transcriptional regulators.

How is the Rnf complex central to *F. nucleatum* metabolism and virulence? We propose that the *F. nucleatum* Rnf complex promotes metabolism of many amino acids, via the oxidation of reduced ferredoxin to the reduction of NAD+ that generates an electrochemical ion gradient across the cytoplasmic membrane, leading to the formation of several key metabolites and ATP (Fig. 6). As such, genetic disruption of the Rnf complex causes pleiotropic defects that negatively impact the pathophysiology of *F. nucleatum*. Given the wide conservation of the Rnf complex in many pathogens and its absence in eukaryotes, this multi-subunit respiratory enzyme may serve as an important target for broad anti-infective therapeutic strategies.

## MATERIALS AND METHODS

### Bacterial strains, plasmids, and media

The bacterial strains and plasmids used in this study are listed in *Supplemental Information* (*SI*) Table S1. *F. nucleatum* cells were grown in tryptic soy broth (TSB) supplemented with 1% Bacto^TM^ peptone and 0.25% fresh cysteine (TSPC) or on TSPC agar plates in an anaerobic chamber (10% CO_2_, 10% H_2_, and 80% N_2_). Heart infusion broth (HIB) or agar (HIA) was used to culture *A. oris* and supplemented with 0.5% glucose to grow *S. oralis* and *S. gordonii. E. coli* strains were grown in Luria broth (LB). All bacterial strains, except *F. nucleatum*, were cultured in a 5% CO_2_ incubator. When required, chloramphenicol or thiamphenicol were added into the medium at a concentration of 15 µg/ml and 5 µg/ml, respectively. All reagents were purchased from Sigma-Aldrich unless noted otherwise.

### Plasmid construction

**(i)** To generate pRnfC, the primer set com-rnfC-F/R (*SI* Table S2) was used to amplify the *rnfC* coding sequence and its promoter region from chromosomal DNA of *F. nucleatum* ATCC 23726, while appending the restriction sites KpnI/NdeI to the amplicon. The PCR product was digested with KpnI and NdeI restriction enzymes and cloned into pCWU6 (*SI* Table S2) precut with the same enzymes. **(ii)** To generate pRnfD, a segment encompassing the *rnfC* promoter region and the *rnfD* coding region was PCR-amplified from chromosomal DNA of *F. nucleatum* Δ*rnfC*, using the primer set com-rnfD-F/R (SI Table S2), while appending NdeI and XhoI restriction sites to the amplicon. The PCR product was digested with NdeI and XhoI restriction enzymes and cloned into pCWU6 precut with the same enzymes. (**iii**) To generate pMCSG7-RnfC, the primer set, LIC-RnfC-F/R, was used to PCR-amplify the *rnfC* coding sequence from chromosomal DNA of *F. nucleatum* ATCC 23726, while appending adapter sequences for subsequent ligation-independent cloning (LIC) into pMCSG7 as previously reported (47). All generated vectors were subjected to DNA sequence to confirm cloned sequences.

### Gene deletion in *F. nucleatum*

Generation of non-polar, in-frame deletion mutants, Δ*rnfC* and Δ*rnfD*, was performed according to our published protocol (24, 26, 37), with primers for generation of deletion constructs listed in *SI* Table S2.

### qRT-PCR

Overnight-grown (∼ 17 h) fusobacterial cultures were normalized to OD_600_ of ∼2.0, and cells were harvested by centrifugation for RNA extraction using RNeasy Mini Kits (Qiagen) according to the manufacturer’s instructions, as described previously (25). Purified RNA, free from DNA by treatment with DNase I (Qiagen), was used for cDNA synthesis using iScript^TM^ RT supermix (Bio-Rad) based on the manufacturers protocol. Real-time PCR reactions were prepared using the SYBR® Green PCR Master Mix with the appropriate primers (Table S2), and analysis was performed using the CFX96 Real-Time System (Bio-Rad). The ΔΔC_T_ method was used to calculate changes in gene expression between samples. Briefly, ΔΔC_T_ = ΔC_T1_ - ΔC_T2_, where ΔC_T_ = C_T_ (target) - C_T_ (housekeeping gene). Fold changes were calculated as log_10_(2^ΔΔCt^). The *16S* rRNA gene was used as reference. Reactions without reverse transcriptase were used as control to assess genomic DNA contamination.

### Biofilm assay

Stationary-phase cells of fusobacterial strains normalized to an OD_600_ of ∼0.6 in fresh TSPC were seeded into flat-bottom, multi-well plates (Greiner Bio-One) and grown anaerobically for 48 h at 37°C. Obtained biofilms were washed gently in phosphate-buffered saline (PBS) and dried before being stained with 0.2 ml of 1% crystal violet (w/v) solution for 10 min at room temperature. Biofilms were gently washed with sterile water before drying and imaging. For quantification, dried biofilms were de-stained with acetic acid (30% v/v) for 10 min at room temperature, followed by measurement at OD_590_. Results were obtained from at least three independent experiments performed in triplicate. Statistical analysis was performed by GraphPad Prism 9.0.

### Bacterial coaggregation assays

Fusobacterial interaction with bacterial partner strains was assessed using a previously published coaggregation assay (24), with some modifications. Briefly, overnight cultures of *F. nucleatum* and partner strains were harvested by centrifugation and washed twice in coaggregation buffer [0.02 M tris-buffered saline, 150 mM NaCl, 0.1 mM CaCl_2_]. Fusobacterial cells were normalized to an OD_600_ of ∼0.4 in coaggregation buffer without or with 15 mM L-lysine, when necessary, whereas partner strains were normalized to an OD_600_ of 2.0 in coaggregation buffer. 0.25 ml aliquots of fusobacterial and partner strains were mixed in multi-well plates (GenClone), and coaggregation was imaged. For the experiments using unwashed fusobacterial cells, a similar procedure was used, except that fusobacterial cultures were directly used without washing. Results were obtained from at least three independent experiments performed in triplicate.

### H_2_S detection

Detection of H_2_S was performed according to a published protocol with slight modification (25). Briefly, overnight cultures of fusobacterial cells were normalized to an OD_600_ of ∼0.6 in TSPC, and 0.1 ml of normalized cell suspension was mixed with 0.1 ml of bismuth solution [0.4 M triethanolamine pH 8.0, 10 mM bismuth chloride, 20 mM pyridoxal 5-phosphate monohydride, 20 mM EDTA, 40 mM L-cysteine] and incubated at 37°C anaerobic for 1 h, at which point images were taken and H_2_S was quantified by OD_405_. The results were presented as average of at least three independent experiments performed in triplicate.

### Measurement of lysine levels by liquid chromatography-mass spectrometry

The relative lysine levels from the culture supernatant of various fusobacterial strains grown to a stationary phase were determined at the Metabolomics Core at Baylor College of Medicine, according to our previous publication (24). Significance analysis was performed by GraphPad Prism.

### ATP quantification assay

Mid-log phase fusobacterial cells (OD_600_ of ∼ 0.5) were harvested by centrifugation and combined in equal volumes (0.1 ml) with the BacTiter-Glo Reagent (Promega) in multi-well plates (Cellstar). The plates were incubated at room temperature in dark for ∼2 min prior to measurement of relative luminescent units (RLU’s) using a plate reader (Tecan). ATP concentrations were determined from an ATP standard curve with ATP concentrations ranging from 1 to 1000 nM. The results, presented as pmol/10^6^ CFU, were obtained from at least three independent experiments performed in triplicate.

### Western blotting

Expression of fusobacterial proteins was analyzed by immunoblotting with antibodies against RadD (α-RadD; 1:5000), MegL (α-MegL; 1:5000), FtsX (α-FtsX; 1:2000) and RnfC (α-RnfC; 1:3000). Generation of the first 3 antibodies was described elsewhere (24-26). For α-RnfC, the procedure was followed according to a previously published protocol (26). Briefly, *E. coli* BL21 (DE3) harboring pMCSG7-RnfC was used to purify recombinant protein RnfC by affinity chromatography. The purified protein was used for antibody production (Cocalico Biologicals, Inc.). For immunoblotting, overnight fusobacterial cultures were normalized to an OD_600_ of 1.0, and 1-ml aliquots of bacterial cultures were taken for trichloroacetic acid (TCA) precipitation and acetone-wash, as previously reported (24). Protein samples obtained were suspended in SDS-containing sample buffer, separated by SDS-PAGE using 4-15% Tris-Glycine gradient gels (RadD and MegL) or a 12% Tris-Glycine gel (RnfC), and immunoblotted by specific antibodies.

### Electron microscopy

Electron microscopy was performed according to previously published protocols (24, 26). Briefly, overnight cultures of fusobacteria were harvested by centrifugation and re-suspended in PBS supplemented with 0.1 M NaCl. A drop of bacterial suspension was added to carbon-coated grids, stained with 1% uranyl acetate, and washed prior to imaging with an electron microscope (JEOL 1200).

### Immunofluorescence microscopy

Overnight-grown fusobacterial cells were harvested by centrifugation and washed twice in PBS before normalizing to an OD_600_ of 0.5 in PBS. 0.2-ml aliquots of cell suspension were used to coat circular glass coverslips placed in a 24-well plate for 20 min at room temperature. Cells were fixed with 2.5% formaldehyde (in PBS) for 20 min, washed with PBS, and blocked with 3% w/v bovine serum albumin (BSA) for 1 h. Cells were incubated with α-RadD (1:200) for 1 h and then AlexaFluor 488 goat anti-rabbit IgG for another hour, followed by washing in PBS three times in the dark. Cells were analyzed by a fluorescence microscope (Keyence BZ-X800).

### Targeted metabolic analysis

Parent and Δ*rnfC* mutant cells were harvested by centrifugation from mid-log phase cultures in triplicate and normalized to an OD_600_ of 0.5 before drying in a speed vacuum concentrator (Thermo Scientific). Dried pellet samples were subjected to metabolomics analysis at the UC Riverside Metabolomics Core Facility as previously described with some modifications (48). In brief, 10 mg of dry pellet was extracted with 100 µl/mg of monophasic extraction solvent (30:30:20:20/acetonitrile, methanol, isopropanol and water) by sonication on ice for 5 min, followed by 90-min vortexing at 4°C. Cell-free supernatants were collected by centrifugation (3000 g) at 4°C for 30 min, and 1-ml aliquots were analyzed by a TQ-XS triple quadrupole mass spectrometer (Waters) coupled to an I-class UPLC system (Waters), with separations performed using a ZIC-pHILIC column (2.1 x 150 mm, 5 µM) (EMD Millipore). The mobile phases were (A) water supplemented with 15 mM ammonium bicarbonate titrated to a pH of 9.6 with ammonium hydroxide and (B) acetonitrile. The flow rate was 200 µL/min and the columns were held at 50°C. The injection volume was 2 µL. The gradient was as follows: 0 min, 90% B; 1.5 min, 90% B; 16 min, 20% B; 18 min, 20% B; 20 min, 90% B; 28 min, 90% B. The MS was operated in multiple reaction monitoring mode (MRM). Source and desolvation temperatures were 150° C and 600° C, respectively. Desolvation gas was set to 1100 L/h and cone gas to 150 L/h. Collision gas was set to 0.15 mL/min. All gases were nitrogen except the collision gas of argon. Capillary voltage was 1 kV in positive ion mode and 2 kV in negative ion mode. A quality control (QC) sample, generated by pooling equal aliquots of each sample, was analyzed periodically to monitor system stability and performance. Samples were analyzed in random order. Statistical analysis was performed relative to the parent strain and determined by a Student’s *t* test, using GraphPad Prism 9.0.

Raw mass-spectral data for all 81 identified metabolites with corresponding KEGG identifiers above for *F. nucleatum* ATCC 23726 were used to perform quantitative enrichment and pathway analysis using MetaboAnalyst 5.0 (49, 50). Statistical analysis was performed relative to the parent strain and determined by the Global test of MetaboAnalyst 5.0.

### Mouse model of preterm birth

The virulence potential of Δ*rnfC* was evaluated using a published mouse model of preterm birth (25, 37). Briefly, groups of five CF-1 (Charles River Laboratories) pregnant mice were infected with ∼ 5 x 10^7^ CFU of the parent or Δ*rnfC* strain at day 16 or 17 of gestation via tail vein injection. Pup survival was recorded for the next 7 days. Statistical analysis was determined via the Mantel-Cox text, using GraphPad Prism 9.0, and specified in corresponding figure legends.

All animal procedures were approved by the UCLA Animal Research Committee.

## ACKNOWLEDGMENTS

We thank Amancio De Souza (Metabolomics Core Facility, UCR) for technical assistance and our lab members for their discussion and critical review of the manuscript. Research reported in this publication was supported by the National Institute of Dental & Craniofacial Research (NIDCR) of the National Institutes of Health under Award Numbers DE030895 (to C.W.) and DE026758 (to H.T.-T). T.A.B. was supported by the UCLA Dentist-Scientist and Oral Health-Researcher Training Program, NIDCR Grant T90DE030860. D.F. was supported by the Ruth L. Kirschstein National Research Service Award (T32AI007323). The content is solely the responsibility of the authors and does not necessarily represent the official views of the National Institutes of Health.

## AUTHOR CONTRIBUTIONS

T.A.B. and H.T.-T. designed research; T.A.B., C.W., Y.-W.C., D.F., Y.C., M.I.C., and T.T.L. performed research; T.A.B, C.W., Y.-W.C., and A.D. analyzed data; and T.A.B., A.D., and H.T.-T. wrote the paper.

## DECLARATION OF INTERESTS

The authors declare no competing interests.

## Notes

### Competing Interest Statement

The authors have declared no competing interest.

### Summary of Updates

The title, abstract, and Importance have been revised; Figure 6 revised; Supplemental files updated

