## Supporting Information for "The respiratory enzyme complex Rnf is vital for metabolic adaptation and virulence in *Fusobacterium nucleatum*"

[a]*Molecular Biology Institute, University of California, Los Angeles, California, USA;* [b]*Department of Microbiology & Molecular Genetics, University of Texas McGovern Medical School, Houston, Texas, USA;* [c]*Division of Oral & Systemic Health Sciences,, School of Dentistry, University of California, Los Angeles, California, USA;* [d]*Department of Medicine, Neag Comprehensive Cancer Center, University of Connecticut Health Center, Farmington, CT, USA;* [e]*Department of Microbiology, Immunology & Molecular Genetics, University of California, Los Angeles, Los Angeles, CA, USA.*

Running Title: Rnf complex in bacterial virulence

Keywords: Rnf complex, metabolism, hydrogen sulfide, coaggregation, preterm birth, virulence

### Supporting Figures

**Figure S1: Expression analysis of RnfC by western blotting. (A)** Protein samples in the whole-cell lysates prepared from the parent and Tn5 mutant strains were immunoblotted with antibodies against RnfC (α-RnfC). **(B)** Protein samples obtained from whole-cell lysates of the parent, its isogenic *rnfC* deletion mutant Δ*rnfC*, and *rnfC*-complemented Δ*rnfC*/pRnfC strains were immunoblotted with α-RnfC. An antibody against the membrane-bound cell-division protein FtsX was used as a loading control. All results were obtained from three independent experiments performed in triplicate.

**Figure S2: RnfC is dispensable for surface display of RadD. (A)** Overnight-grown cells of indicated fusobacterial strains were first stained an antibody against RadD (α-RadD) and then an Alexa Fluor 488-secondary antibody. Surface display of RadD was analyzed by fluorescence microscopy. **(B)** Expression of RadD was analyzed by immunoblotting of protein samples in the whole-cell lysates obtained from the indicated strains. A Coomassie-stained band (*) was used as loading control. The results presented are representative of three independent experiments performed in triplicate.

**Figure S3: Deletion of *rnfC* significantly reduces expression of genes coding for enzymes involved in $H_2S$ production and lysine catabolism.** Normalized overnight cultures of the parent, in Δ*rnfC*, and Δ*rnfC*/pRnfC strains were used to isolate total RNA. The expression levels of *megL*, *cysK1*, *cysK2*, *kamA*, *kamD*, and *radD* in these strains were determined by qRT-PCR. Results were obtained from three independent experiments performed in triplicate. All qRT-PCR data were normalized to the transcript abundance of 16s rRNA for each sample. Significance was calculated by a student’s t-test; ****, $P < 0.0001$.

**Figure S4: Deletion of *rnfD* causes pleiotropic defects. (A)** Interaction between *S. gordonii* DL1 and indicated fusobacterial strains was determined by a coaggregation assay, with fusobacterial cells washed or unwashed prior to mixing with *gordonii*. A *radD* mutant was used as a negative control. **(B)** Bacterial growth of indicated strains was monitored by optical density at 600 nm over 24 h. **(C)** Biofilms of indicated strains were cultivated for 48 h of anaerobic incubation. Quantification of biofilm production was determined by 1% crystal violet staining. **(D)** Normalized overnight cultures of indicated fusobacterial strains were used to determine hydrogen sulfide production by a bismuth assay. All results were obtained from three independent experiments performed in triplicate, and significance calculated by a student's t-test; ****, $P < 0.0001$.

**Figure S5: Deletion of *rnfC* alters methionine and cysteine metabolism. (A)** Using the MetaboAnalyst 5.0 web-based software, pathway analysis was performed on differentially expressed metabolites between the parent and Δ*rnfC* strain (see Fig. 4C). Shown is the generated pathway from the "Cys/Met metabolism" node. The level of significance for a metabolite abundantly detected in these two strains is indicated by yellow to red (P value ranging from 0.3 to 0.002). KEGG identification numbers of metabolites are highlighted in light blue. **(B-C)** Shown are relative concentrations of methionine (B) and S-adenosyl-L-homocysteine (C). Statistical significance was calculated using the Global test.

### Supporting Tables

#### Table S1: Bacterial strains and plasmids used in this study

| Strains & Plasmids | Description | Reference |
|---|---|---|
| *Strain* | | |
| *F. nucleatum* ATCC 23726 | Type strain | (1) |
| *F. nucleatum* CW1 | Derivative of 23726; lacking *galK* (Δ*galK*) | (1) |
| *F. nucleatum radD*::Tn5 | Derivative of ATCC 23726 with Tn5 insertion into *radD* at position 5615 (10386) | (2) |
| *F. nucleatum rnfA::*Tn5 | Derivative of ATCC 23726 with Tn5 insertion into *rnfA* at position 511 (585) | (1) |
| *F. nucleatum rnfB::*Tn5 | Derivative of ATCC 23726 with Tn5 insertion into *rnfB* at position 222 (1158) | (2) |
| *F. nucleatum rnfC::*Tn5 | Derivative of ATCC 23726 with Tn5 insertion into *rnfC* at position 222 (1326) | (2) |
| *F. nucleatum* Δ*rnfC* | Isogenic derivative of CW1 lacking *rnfC* | This study |
| *F. nucleatum* Δ*rnfD* | Isogenic derivative of CW1 lacking *rnfD* | This study |
| *F. nucleatum* Δ*megL* | Isogenic derivative of CW1 lacking *megL* | This study |
| *F. nucleatum* Δ*radD* | isogenic derivative of CW1 lacking *radD*) | (1) |
| *F. nucleatum* Δ*kamA* | Isogenic derivative of CW1 lacking *kamA* | (2) |
| *A. oris* MG1 | Type strain | (3) |
| *S. oralis* 34 | RPS positive | (4) |
| *S. gordonii* DL1 | Type strain | (5) |
| *Plasmid* | | |
| pCWU6 | Derivative of pHS30 | (1) |
| pCM-GalK | *C. perfringens* vector expressing *galK* | (1) |
| pMCSG7-RnfC | Recombinant vector expressing His-tagged RnfC | This study |
| pRnfC | Derivative of pCWU6 expressing *rnfC* under the control of a *rpsJ* promoter | This study |
| pRnfD | Derivative of on pCWU6 expressing *rnfD* under the control of a *rpsJ* promoter | This study |
| pGalK--Δ*rnfC* | pCM-galK derivative; *rnfC* deletion vector | This study |
| pGalK--Δ*rnfD* | pCM-galK derivative; *rnfD* deletion vector | This study |
| pGalK--Δ*radD* | pCM-galK derivative; *radD* deletion vector | (2) |
| pGalK--Δ*megL* | pCM-galK derivative; *megL* deletion vector | (6) |
| pGalK--Δ*kamA* | pCM-galK derivative; *kamA* deletion vector | (2) |

**Table S2: Primers used in this study**

| Primer | Sequence[(a)] | Used for |
|---|---|---|
| rnfC-up-F | GGCGGGATCCATGAACTTTGAAGAAATAGATTTTTATATT | pGalK-Δ*rnfC* |
| rnfC-up-R | GGCGGGATCCTTAAAGGAGCTCCTATATGTTGTAAAAG | pGalK-Δ*rnfC* |
| rnfC-dn-F | GGCGGGTACCGTCCTATGGGGCTTGCACCACTTATG | pGalK-Δ*rnfC* |
| rnfC-dn-R | GGCGAAGCTTGCTAGTTGCTTCTGGTAAAACTTCTTTT | pGalK-Δ*rnfC* |
| com-rnfC-F | GGCGGGTACCGGATAGTAGAAGTGCATTTAAAGATT | pRnfC |
| com-rnfC-R | GGCGGGATCCCTACTTTTTCTTAGCTCTTAATTTAG | pRnfC |
| LIC-RnfC-F | TACTTCCAATCCAATGCAATGAAAGGAGTGTTT | pMCSG7-RnfC |
| LIC-RnfC-R | TTATCCACTTCCAATGTTACTACTTTTTCTTAGCTC | pMCSG7-RnfC |
| rnfD-up-F | CGCGGATCCAAAGGTATTGTTGGTATAGGAG | pGalK-Δ*rnfD* |
| rnfD-up-R | CCCATCCACTAAACTTAAACAATATGAGGAGCTGGTCCTGT | pGalK-Δ*rnfD* |
| rnfD-dn-F | TGTTTAAGTTTAGTGGATGGGTTTGCATTGGGATTAGGAGTTT | pGalK-Δ*rnfD* |
| rnfD-dn-R | CGCGTCGACAAATAATCCTAATACCTTATATAAG | pGalK-Δ*rnfD* |
| com-rnfD-F | GGGAATTCCATATGGTTTAGGAAATCCGGGCAAA | pRnfD |
| com-rnfD-R | CCGCTCGAGAGCTGCTATTAGACCAAGGA | pRnfD |
| radD-up-F | AAAGTCGACATGGTTTAGTGAAAGATTATTCAAAAT | pGalK-Δ*radD* |
| radD-up-R | AAAGGTACCATTTGCTCCA AAATCTATTT TATCA | pGalK-Δ*radD* |
| radD-dn-F | AAAGGTACCTCATCATCACCAATATTTAAGTCATTAG | pGalK-Δ*radD* |
| radD-dn-R | AAAGAGCTCCATAAATATCCTCAAAATATGAGTG | pGalK-Δ*radD* |
| kamA-up-F | GGCGTGAGCTCCAGAGATAGAAGTTTTTGATAAGGGTA | pGalK-Δ*kamA* |
| kamA-up-R | GGCGAGGTACCGTTTACCTTTCTACTACCATACCATAAT | pGalK-Δ*kamA* |
| kamA-dn-F | GGCGAGGTACCGGTACCAAAATAAAAAATGTTAGATAC | pGalK-Δ*kamA* |
| kamA-dn-R | GGCGAGTCGACAAGTAGCAATTTTTTCATTATTAGGAT | pGalK-Δ*kamA* |
| megL-up-F | CGCGGATCCGACATTCTCTTGAATTATAAAAAAAATCTG | pGalK-Δ*megL* |
| megL-up-R | AAAACTGCAGCCATTATAGATTTCTTTCCCATAACC | pGalK-Δ*megL* |
| megL-dn-F | TAACTTTACTCATTTGTCTTAATTCCTTAC | pGalK-Δ*megL* |
| RT-megL-F | CACAAGACTAGGCAATCCTACA | RT-PCR *megL* |
| RT-megL-R | GCTCCCATACCAGATGACATAG | RT-PCR *megL* |
| RT-cysK1-F | AACAGGGACAGGAGGTAGTT | RT-PCR *cysK1* |
| RT-cysK1-R | AGATGAAGCAGGCTCAACAG | RT-PCR *cysK1* |
| RT-cysK2-F | GCTACAAGTGGAAACACAGGA | RT-PCR *cysK2* |
| RT-cysK2-R | TCACTCATCCAATCTGGCATATAA | RT-PCR *cysK2* |
| RT-kamA-F | TCTCAATGGCAACTGGATTCTC | RT-PCR *kamA* |
| RT-kamA-R | TGCAGCATGGTCAACTGTATAA | RT-PCR *kamA* |
| RT-kamD-F | GTGCTGATGTTGTTGCAGTTAT | RT-PCR *kamD* |
| RT-kamD-R | TTCTTGTGTTGCCATTGTTCC | RT-PCR *kamD* |
| RT-radD-F | GCAGCAGCACCAACAATAAAT | RT-PCR *radD* |
| RT-radD-R | GGTGCTTCAGGAGGTGTTATC | RT-PCR *radD* |
| RT-16s-F | GGTTAAGTCCCGCAACGA | RT-PCR *16s* |

| | | |
|---|---|---|
| RT-16s-R | CATCCCCACCTTCCTCCTAC | RT-PCR *16s* |

[a] Underlined are restriction site sequences.

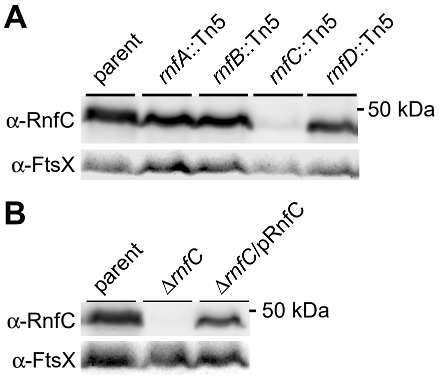


Figure S1: Britton et al.

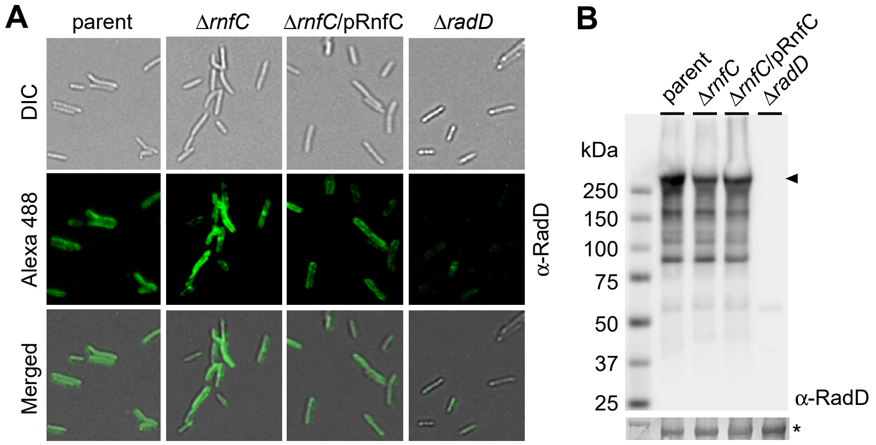


Figure S2: Britton et al.

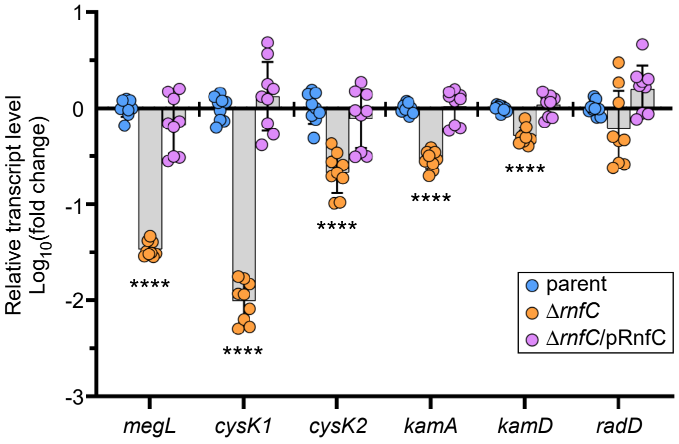


Figure S3: Britton et al.

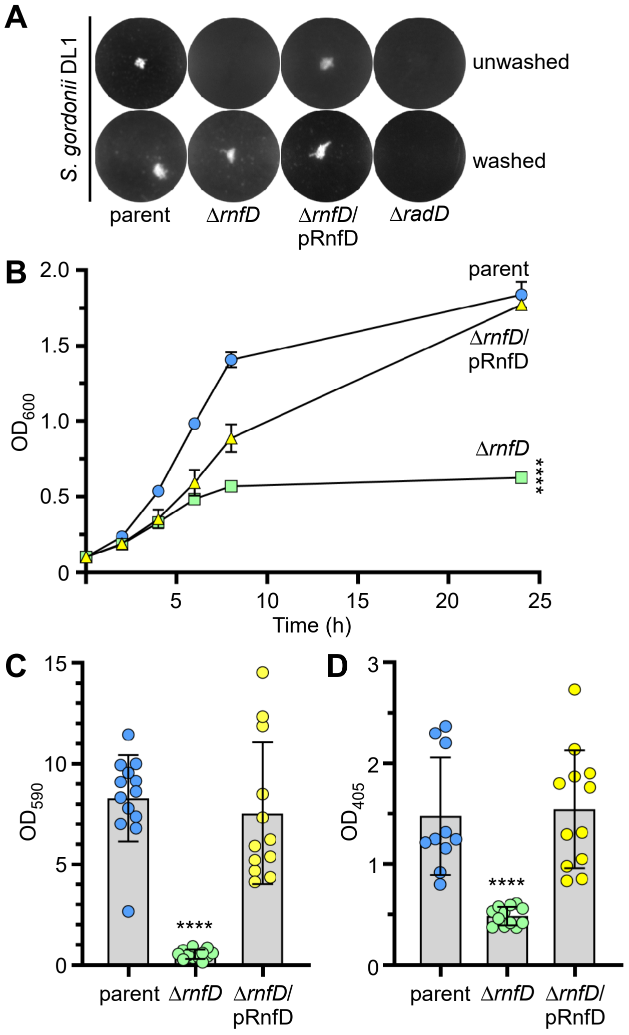


Figure S4: Britton et al.

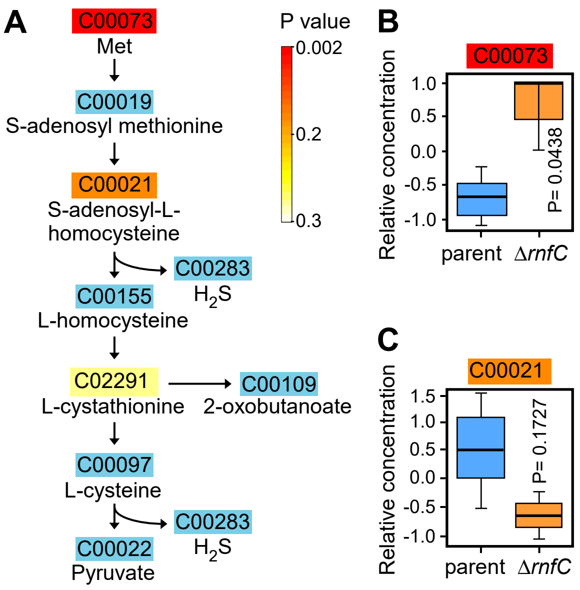


Figure S5: Britton et al.
